# An Efficient and Cost-effective Purification Methodology for SaCas9 Nuclease

**DOI:** 10.1101/2021.06.08.447622

**Authors:** Allen C.T. Teng, Marjan Tavassoli, Suja Shrestha, Kyle Lindsay, Evgueni Ivakine, Ronald Cohn, J. Paul Santerre, Anthony O. Gramolini

## Abstract

With an ever-increasing demand for laboratory-grade Cas9 proteins by many groups advancing the use of CRISPR technology, a more efficient and scalable process for generating the proteins, coupled with rapid purification methods is in urgent demand. Here, we introduce a modified methodology for rapid purification of active SaCas9 protein within 24 hours. The product has over 90% protein purity. The simplicity and cost-effectiveness of such methodology will enable general labs to produce a sizable amount of Cas9 proteins, further accelerating the advancement of CRISPR/Cas9-based research.

## Introduction

Precise editing of genomic DNA remained challenging until the advent of the clustered regularly interspaced short palindromic repeats (CRISPR)-associated protein (Cas) system. CRISPR/Cas was first identified as an adaptive immune system in bacteria and archaea as a defense against invading viruses and plasmids ^1^. Different types of Cas proteins can target double stranded DNA, single stranded DNA, or RNA ^2–6^. One best characterized Cas subtype is Cas9, which has now been routinely used for modifying genomic DNA of model organisms in laboratories. Of the many identified Cas9 proteins, Cas9 from *Staphylococcus aureus* (SaCas9) has emerged as a preferred genetic editing tool, because of its smaller size (1053 amino acids) allowing for an efficient encapsulation into transfecting complexes or transducing viral DNA ^7,8^. For example, SaCas9 has been extensively used as a genome editing tool for developing an array of therapeutic strategies investigating human inherited diseases in animal models ^2,9^. However, with an increasing demand, but currently time consuming and high cost-associated methods for generating purified SaCas9, acquiring this protein remains a challenge for many laboratories that are not adequately equipped for protein purification. Here, we report an advancement of the purification methods for SaCas9 from bacterial cells. A major advantage of this methodology is associated with achieving over 90% purity, at large batch sizes (concentrations at 1 mg/L) within a day. Purified SaCas9 can be directly used for *in vitro* applications. The new methodology is superior to the majority of conventional approaches, which depends on French press, high-frequency sonicator, or fast protein liquid chromatography ^10,11^.

## Materials and Methods

### Materials

A glycerol stock of XJb autolysis *E. coli* cells (ZYMO Research, Irvine, CA) containing the His_8_-TEV-SaCas9 (MN_548085.1) expression plasmid. LB broth, LB agar, carbenicillin, Isopropyl-b-D-thiogalactopyranoside, L-arabinose, MgCl_2_, dithiothreitol, imidazole, PMSF are from Bioshop (Toronto, Canada). HEPES, KCl, NaCl, glycerol, MgCl_2_, 2,2,2-Trichloroethanol, and Amicon Ultra-15 centrifugal filter units are from Sigma-Aldrich. Universal Nuclease (Pierce). HisTrap high performance and HiTrap SP HP columns are from Cytiva.

## Methods

### Overexpression of His_8_-TEV-SaCas9 proteins in *E. coli*

XJb *E. coli* cells (ZYMO research, Irvine, CA) containing His8-TEV-SaCas9 expression plasmids were grown in a 4 L LB broth (Bioshop, Toronto, Canada) containing 100 μg/mL carbenicillin (Bioshop, Toronto, Canada) at 37°C/250 rpm until optical density 600 nm (O.D.600) reached 0.6~0.8 at 37°C. Induced protein expression with 0.5 mM IPTG (final concentration) and 3 mM L-arabinose solution (final concentration) overnight at 180 rpm, 18°C. Next morning, bacteria pellet was collected by centrifugation at 6,000x*g* and suspended 1 L of bacteria pellet with 40 mL buffer A supplemented with 1 mM PMSF. Homogenized bacteria can be stored at - 80°C until protein purification.

### Ni^2+^-NTA column purification of His_8_-SaCas9 proteins

Cells were lyzed in a 37°C water bath for 20 minutes and ribonucleic acids were digested with 0.75 μL of universal ribonuclease (Pierce, 88701) at 37°C for 1 hour. Insoluble fraction was removed by spinning protein lysate at 15,000 *xg* for an hour and then soluble proteins were filtered through a 0.22 μm filter. Ni^2+^-NTA column connected to a syringe pump was set up as shown in Figure 1. Equilibrated column with 25 mL ddH_2_O, 25 mL solution B (20 mM HEPES; pH 7.5, 300 mM NaCl, 250 mM Imidazole, 0.5 mM DTT), and 25 mL solution A (20 mM HEPES, pH7.5, 300 mM NaCl, 25 mM Imidazole) at the rate of 4 mL/min. Applied filtered supernatant into the Ni^2+^-NTA column at the rate of 2 mL/min. Washed column with 30 mL washing buffer (21 mL solution A + 9 mL solution B, 30% solution B). Eluted His_8_-TEV-SaCas9 proteins in 15 mL elution buffer (100% solution B). Ran 40 μL eluted proteins on a SDS-PAGE gel for protein visualization.

**Figure 1.**
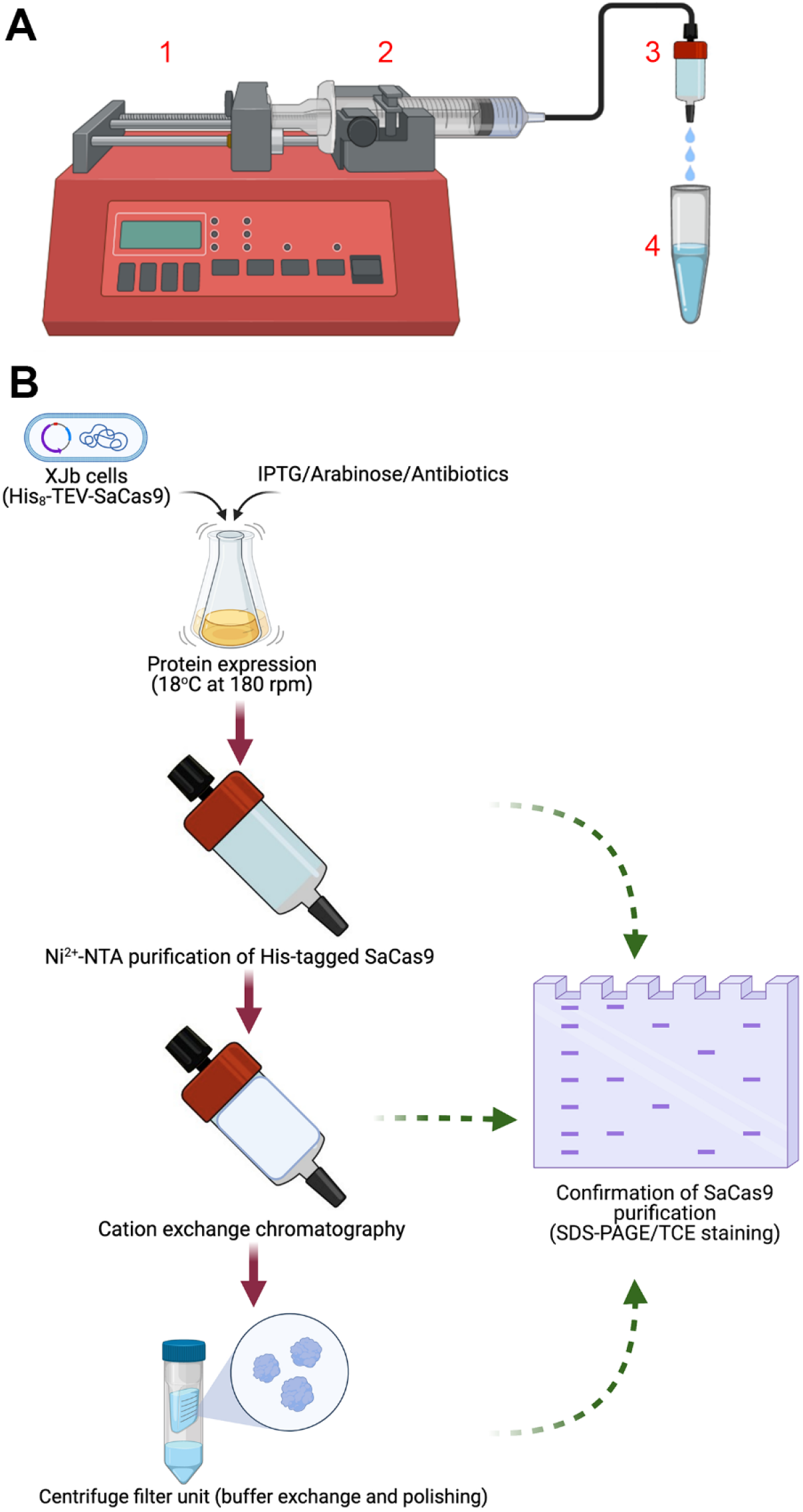
Methodology illustration for His_8_-TEV-SaCas9 purification. (**A**) Apparatus setup for protein purification. A 50-mL syringe (2) mounted on a syringe pump (1) is connected to a pre-packed column (3). Loading of protein samples and buffer application can be swiftly adapted by changing different syringes. Eluted samples can be collected in tubes (4). (**B**) The summary of SaCas9 purification procedure introduced in this study. SaCas9 expression is induced overnight in XJb cells. Protein purification is performed by Ni^2+^-NTA and CIEX chromatography as illustrated in **Fig. 1A**. Final protein is concentrated and buffer exchanged in a centrifugal filter unit. Confirmation of protein purity is monitored by TCE staining of SDS-PAGE gels.

### Cation exchange chromatography purification of His_8_-SaCas9 proteins

Diluted eluted His_8_-TEV-SaCas9 proteins by 3-fold with 30 mL buffer C (20 mM HEPES; pH 7.5, 200 mM KCl, 10 mM MgCl_2_, 0.5 mM DTT) and filter through a 0.22 μm syringe filter. Set up HiTrap SP HP column as shown in Figure 1A. Equilibrated column with 25 mL ddH_2_O, 25 mL solution D (20 mM HEPES; pH 7.5, 1 M KCl, 10 mM MgCl_2_, 0.5 mM DTT), and 50 mL solution C at the rate of 4 mL/min sequentially. Applied filtered sample to the HisTrap column at the rate of 2 mL/min. Washed column with 30 mL washing buffer (24 mL solution C + 6 mL solution D, 20% solution D). Eluted His_8_-TEV-SaCas9 proteins in 20 mL elution buffer (14 mL buffer C + 6 mL buffer D, 30% solution D). Ran 40 μL eluted proteins on a SDS-PAGE gel for protein visualization.

### Buffer exchange and His8-TEV-SaCas9 protein concentration

Concentrated eluted His8-TEV-SaCas9 proteins via a centrifugal filter unit (100 kDa cutoff, Millipore) at 1,000 x*g* for 5 minutes. Performed buffer exchange with 1 mL SaCas9 storage buffer (10 mM Tris-HCl; pH 7.4, 300 mM NaCl, 0.1 mM EDTA, 1 mM DTT, and 50% Glycerol) in the centrifugal filter. Repeat this process 5 times. Transferred SaCas9 proteins into a 1 mL Eppendorf tube and determine protein concentration via Bradford assays. Ran 40 μL eluted proteins on a SDS-PAGE gel for protein visualization.

### Determine SaCas9 enzymatic activity

gRNA and template DNA are prepared as the following. Double stranded DNA (dsDNA) encoding for a sgRNA targeting intron 55 (in55) of the mouse *Dmd* gene (ATG AAA CCA TGG CAA GTA AG) was cloned using BsaI directional cloning into pX601 vector. Briefly, pX601 was digested with BsaI and dephosphorylated with shrimp alkaline phosphatase (New England Biolabs, M0371L). dsDNA in55 guide oligos were designed with 5’overhangs complimentary to the sticky ends produced by BsaI digestion, phosphorylated with T4 polynucleotide kinase (New England Biolabs, M0201L) and ligated into digested pX601. Accurate cloning of the guide was confirmed by Sanger sequencing. In55 guide encoding DNA was amplified from the cloned pX601-in55 vector with Q5 high-fidelity polymerase (New England Biolabs, M0494L), using primers which introduced the minimal T7 promoter sequence upstream of the guide sequence (Fwd: TAA TAC GAC TCA CTA TAG GGA TGA AAC CAT GGC AAG TAA G; Rvs: AAA ATC TCG CCA ACA AGT TG). The amplicon was purified using the QIAquick PCR purification kit (Qiagen, 28104), and sgRNA was *in vitro* transcribed from the amplicon using the MEGAshortscript T7 transcription kit (Thermo Fisher Scientific, AM1354). The sgRNA was purified with the RNEasy Mini Kit (Qiagen, 74104) following the manufacturer’s protocol for RNA cleanup.

SaCas9 was diluted to the final concentration of 500 ng/μL in SaCas9 working buffer (20 mM HEPES, pH 7.4, 150 mM KCl, 10% glycerol, 1 mM DTT), 300 ng/μL gRNA in DNase/RNase-free water, and 60 ng/μL template DNA in DNase/RNase-free water. Mixed together all components as depicted in **Table 2**, vortex, spin down, and incubate at 37°C water bath for 1 hour. Inactivate SaCas9 enzyme at 65°C for 10 minutes. Assessed digestion results on a 1% agarose gel.

**Table 1.**
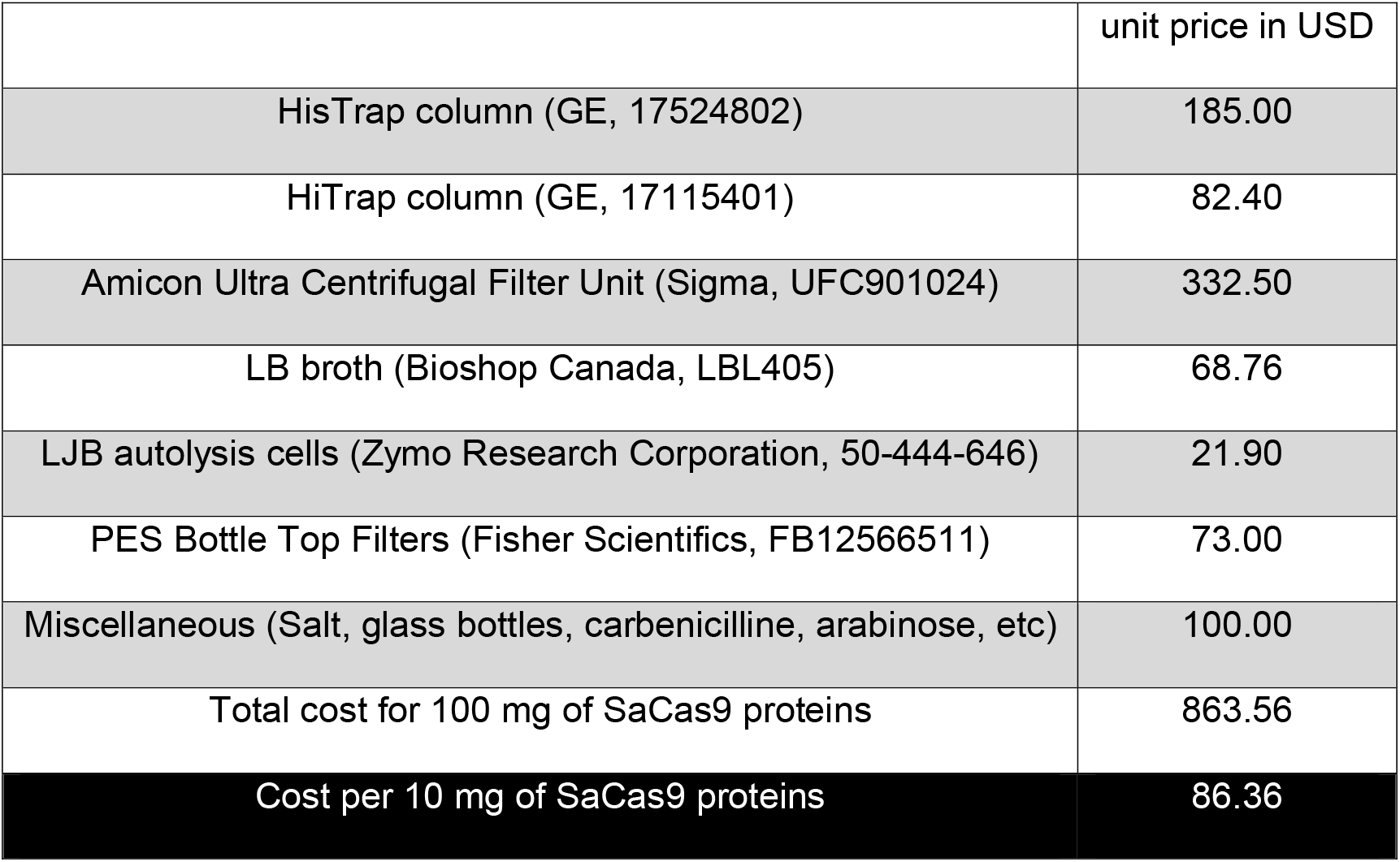

**Table 2.**

|  | SaCas9<br>(500ng/μl) | gRNA<br>(300 ng) | Template<br>DNA (60 ng) | 10X reaction<br>buffer | ddH2O |
| --- | --- | --- | --- | --- | --- |
| Template DNA<br>only |  |  | 1 | 2 | 17 |
| gRNA only |  | 1 |  | 2 | 17 |
| Mock digestion |  | 1 | 1 | 2 | 16 |
| Digestion | 1 | 1 | 1 | 2 | 15 |

## Results

Protein purification is completed by a semi-automated system (**Figure 1A**) and consists of 3 simple steps – (1) an enrichment of His-tagged SaCas9 proteins via a Ni^2+^-NTA column, (2) an intermediate purification via a cation exchange chromatography (CIEX), and (3) and a final buffer exchange via a centrifugal filtration (**Figure 1B**). For the chromatography setup, a 50-mL syringe containing buffer or protein samples is mounted on a syringe pump, which controls the solution flow rate applied to prepacked columns (**Figure 1A**). Eluted protein fractions are monitored by sodium dodecyl sulphate-polyacrylamide gel electrophoresis (SDS-PAGE) with 2, 2, 2-trichloroethanol (TCE) staining. Fractions containing enriched SaCas9 proteins are combined for subsequent fractionations.

Protein purification from bacterial lysates begins with cell lysis ^12–14^. This initial isolation step is commonly achieved using mechanical ^3,4^ or chemical methods ^15^, each with their own limitations. For example, mechanical disruption is efficient in lysing cells, but devices such as a French press or high-frequency sonicator may not be readily available to all laboratories. Chemical lysis of cell walls has a tendency to denature target proteins ^16,17^, further complicating the downstream purification. To overcome these issues, XJb autolysis *E. coli* cells that express λ phage-endolysin for disrupting cell walls were selected as host cells for protein expression. Lysis of *E. coli* containing SaCas9 proteins was achieved by a single freeze-and-thaw process. Successful lysis was accompanied with an increased solution viscosity, which was reduced by a ribonuclease treatment. Next, soluble protein lysate is prepared by centrifugation and degassed by filtration before protein chromatography.

Ni^2+^-NTA chromatography is performed to enrich His_8_-SaCas9. Analyses of SDS-PAGE gels reveals that the majority of His_8_-SaCas9 proteins were eluted in a stepwise fashion between 40% (125 mM) and 100% (275 mM) imidazole solution (**Figure 2A**, asterisk, SaCas9). These fractions are combined and subjected to CIEX chromatography. **Figure 2B** shows that SaCas9 proteins are further fractionated by CIEX chromatography. Specifically, most SaCas9 proteins are eluted stepwise in the presence of 30-40% (500-600 mM KCl) solution D. Keeping K^+^ ion concentrations below 600 mM reduces protein contaminants. Finally, protein concentration and buffer exchange are completed in a centrifugal filtration unit. **Figure 2C** shows that sequentially purified SaCas9 proteins have an average purity of > 90%, based on multiple rounds of protein fractionation. The protein yield from this protocol ranges between 1 mg per liter of bacterial culture. The estimated cost for 10 mg of purified Cas9 proteins is 86.36 USD (**Table 1**).

**Figure 2.**
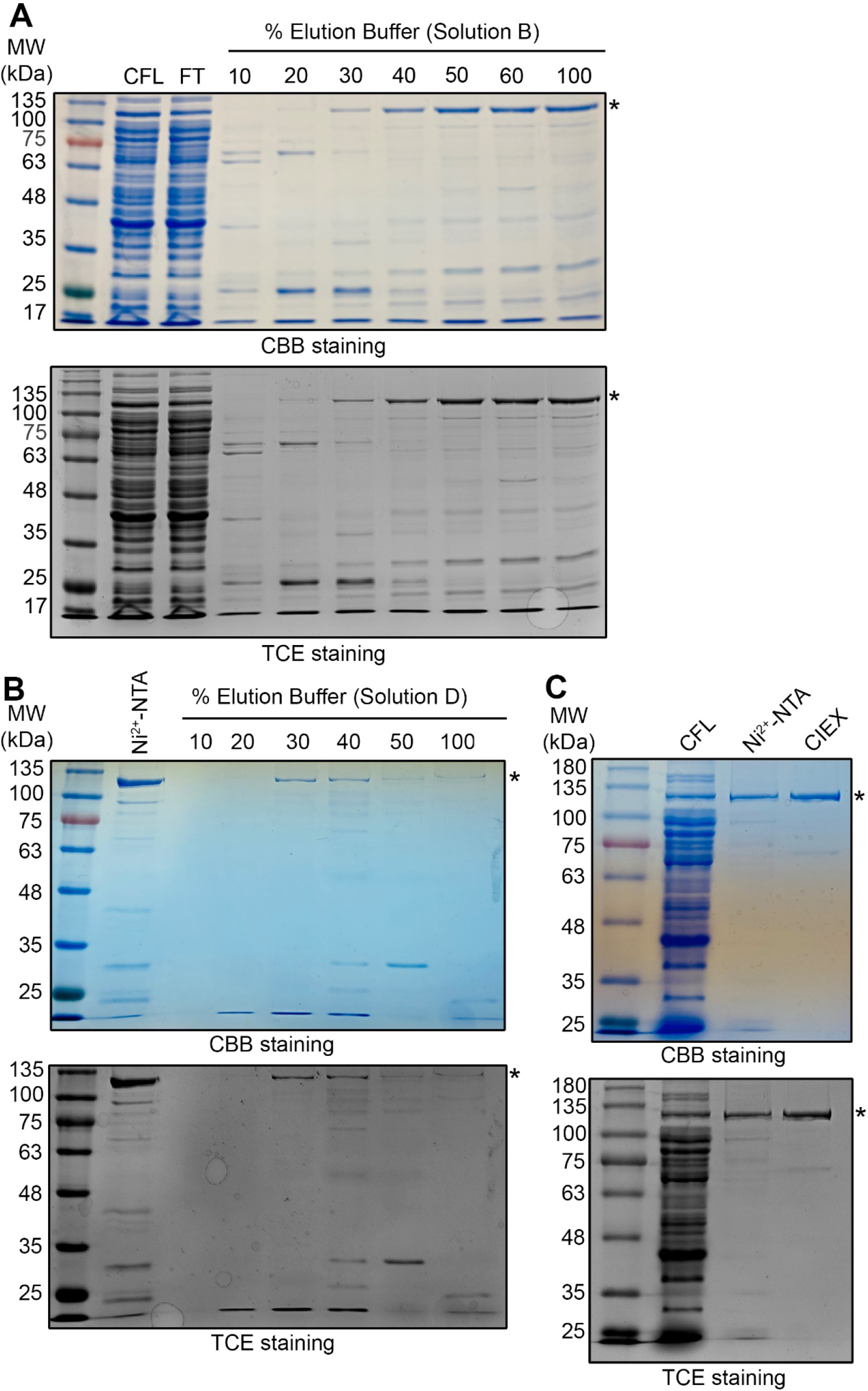
Analyses of Coomassie brilliant blue and TCE stained SDS-PAGE gels for purified His_8_-SaCas9 proteins. (**A**) Affinity purification of His_8_-SaCas9. Soluble His_8_-SaCas9 from bacterial lysates were enriched in a Ni^2+^-NTA column and were eluted in increasing concentration (10-100%) of solution B. Majority of impurity was washed off with 30% solution B and His_8_-SaCas9 proteins were eluted in 15 mL of 100% buffer B. Top panel, CBB staining; Bottom panel, TCE staining. (**B**) Intermediate purification of His_8_-SaCas9 with CIEX. His_8_-SaCas9 proteins from Ni^2+^-NTA-based affinity purification were further fractionated CIEX and proteins were eluted in increasing concentration (10-100%) of buffer D in a stepwise fashion. Majority of SaCas9 proteins were eluted in 30 and 40% fractions and were combined and buffer exchanged in centrifugal filter unit. Top panel, CBB staining; Bottom panel, TCE staining. (**C**) A representative image of sequentially purified His8-SaCas9 proteins. Total proteins from cell-free lysate, Ni^2+^-NTA elution, and centrifugal filtration were visualized on a coomassie brilliant blue-stained SDS-PAGE gel. Top panel, CBB staining; Bottom panel, TCE staining. CFL, cell-free lysate; FT, flow through; *, His8-SaCas9.

Next, we sought to benchmark the purity, production cost, and enzymatic activity of our SaCas9 by comparing it to commercially available sources. **Figure 3A** shows that the purity of this protein is comparable to two commercial suppliers. The production cost associated with this protocol is the lowest - 86.36 USD per 10 mg of proteins (**Table 1**). For determining an activity of purified SaCas9 nucleases, enzymes were mixed with template DNA and gRNA. Digested DNA was visualized on a 1% agarose gel. **Figure 3B** shows that template DNA was completely digested by the *de novo* generated SaCas9 (**Figure 3B**, lane 4) and is comparable to commercially supplied enzymes (**Figure 3B**, lane 3). Together, our results show that purified SaCas9 remains fully active for downstream applications.

**Figure 3.**
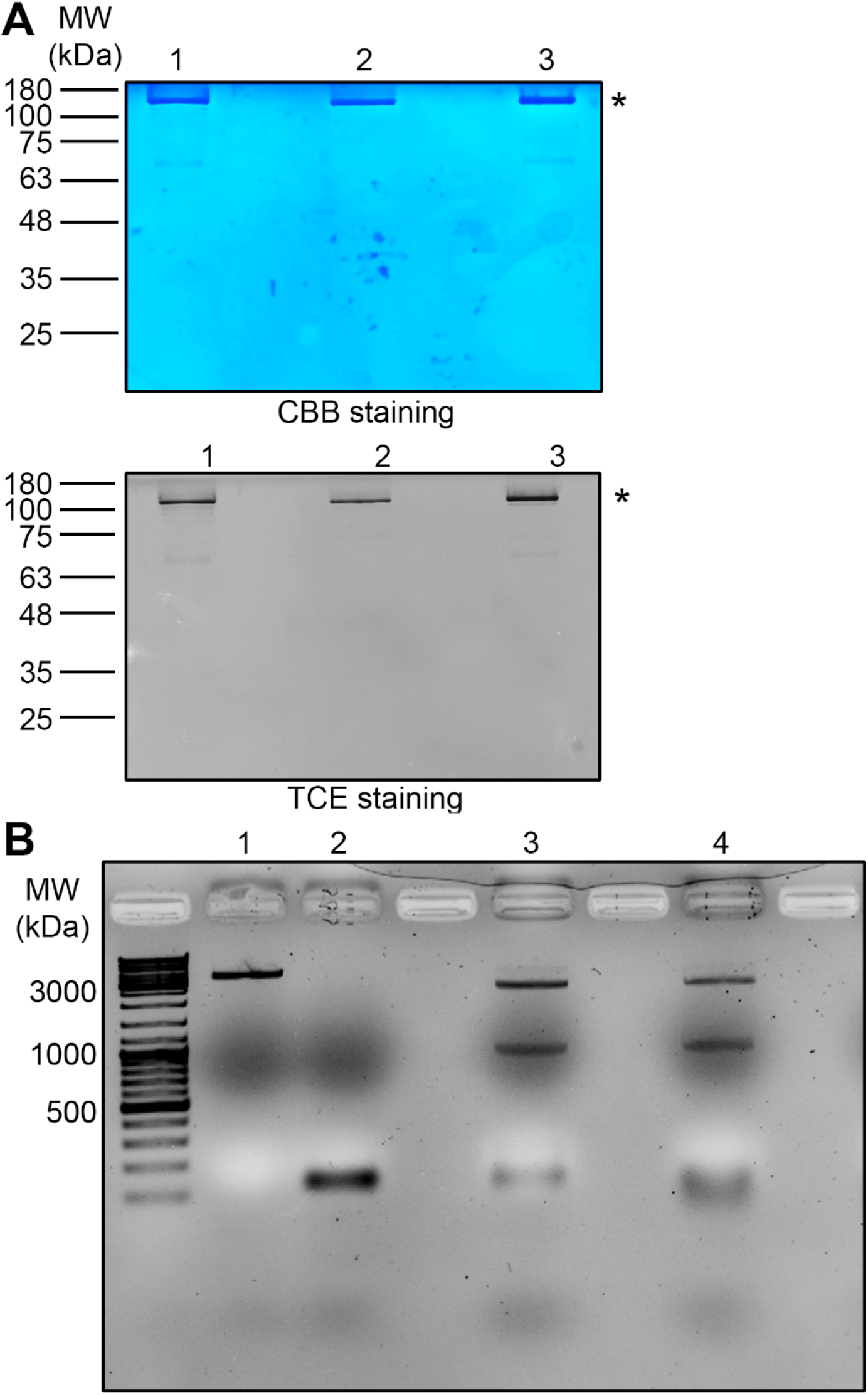
Comparison of homemade SaCas9 to commercial sources. **(A)** Homemade SaCas9 purification procedure achieves 90% purity, which is comparable to commercial suppliers. The purity of *de novo* SaCas9 proteins (1) was visualized side-by-side with 2 other commercial sources (2 & 3) on either CBB or TCE-stained SDS-PAGE gels. The average cost of SaCas9 proteins purified according to the reported method is 8.64 USD/mg. (**B**) Purified SaCas9 efficiently digested double stranded DNA. SaCas9-mediated DNA digestion was analyzed on a 1% agarose gel. A full-length (undigested) template DNA and gRNA were detected in lanes 1 and 2, respectively. A complete digestion of template DNA with commercially supplied SaCas9 was included as a positive control and showed 2 separate fragments (2 and 1 kilobase pairs) in lane 3. A complete digestion of template DNA with homemade SaCas9 was also observed as 2 fragments in lane 4.

## Discussion

In this study, we have demonstrated a modified procedure for purifying SaCas9 from bacterial lysates. The advantages of this method include simplicity and cost-effectiveness. The procedure can be completed within a day by trained personnel and requires only common lab equipment. This process will enable the acceleration of CRISPR/Cas research by providing an easier access to lower cost and highly pure SaCas9 proteins. In addition, different Cas proteins have been purified via Ni^2+^-NTA and CIEX chromatography ^2–4^, thus the reported method can facilitate Cas protein research by further refining the elution buffer concentration for CIEX chromatography. There are, however, two major disclosures associated with this method when compared with fast protein liquid chromatography (FPLC)-based purification. First, the system lacks real-time monitoring for protein elution from chromatography. In modern FPLC, optical density at 280 nm wavelength (O.D._280_) is usually used for keeping track of protein elution. Although such an optical setup is not included in the reported method, SaCas9 can be directly visualized in the TCE-stained SDS-PAGE gels within 30 minutes of chromatography ^18–20^. The second limitation is the absence of a pressure monitor for prepacked columns. Nevertheless, changes of column pressure can be indirectly reflected by the reduced flow rate of solution output. During chromatography, a 20% reduction in flow rate should serve as a sign of increasing column pressure, and the need for an intervention step. Column cleaning and maintenance should be performed as per manufacturer’s manual after three rounds of chromatography, in order to ensure maximal column performance.

## Abbreviations

CRISPR: Clustered Regularly Interspaced Short Palindromic Repeats
SaCas9: Staphylococcus aureus Cas9
CIEX: cation exchange chromatography
SDS-PAGE: sodium dodecyl sulphate-polyacrylamide gel electrophoresis
TCE: 2, 2, 2-trichloroethanol
FPLC: fast protein liquid chromatography
ddH_2_O: double distilled water
IPTG: Isopropyl-b-D-thiogalactopyranoside
dsDNA: Double-stranded DNA

## Acknowledgement

This study was supported by a Medicine by Design New Ideas Fund (MBDNI-201) and a Jesse’s Journey-The Foundation for Gene and Cell Therapy (#507052), and the Translational Biology and Engineering Program (University of Toronto) seed grant.

